# Microglial ablation does not affect opioid-induced hyperalgesia in rodents

**DOI:** 10.1101/2021.04.27.441650

**Authors:** Xin Liu, Bo-Long Liu, Qing Yang, Xiangfu Zhou, Shao-Jun Tang

## Abstract

Opioids are the frontline analgesics in pain management. However, chronic use of opioid analgesics causes paradoxical pain that contributes to the decrease of their efficacy in pain control and the escalation of dose in long-term management of pain. The underling pathogenic mechanism is not well understood. Microglia have been commonly thought to play a critical role in the expression of opioid-induced hyperalgesia (OIH) in animal models. We performed microglial ablation experiments using either a genetic (CD11b-diphtheria toxin receptor transgenic mouse) or pharmacological (colony-stimulating factor 1 receptor inhibitor PLX5622) approaches. Surprisingly, ablating microglia using these specific and effective approaches did not cause detectable impairment in the expression of hyperalgesia induced by morphine. We confirmed this conclusion with behavioral test of mechanical and thermal hyperalgesia, in male and female mice, and with different species (mouse and rat). These findings raise caution about the widely assumed contribution of microglia to the development of OIH.

## Introduction

Morphine and other opioid analgesics are the front-line medicine for managing severe pain[25]. The broad use of opioids for pain management may contribute to the current opioid epidemic, a sharp rise in the incidence of addiction and death by opioid overdose[39]. Clinical data indicate that chronic use of opioids in fact can paradoxically cause a pain state called opioid-induced hyperalgesia (OIH)[10]. The manifestation of OIH is a key factor driving for the dose escalation that is required to effectively control pain, which can eventually lead to potential overdose[3; 11]. Understanding the pathogenic mechanism of OIH is essential for the development of effective strategies to mitigate OIH and improve safety of opioid analgesics.

Although neuronal malplasticity in the pain transmission and modulatory pathways was considered as the main OIH mechanism, emerging evidence indicates a key contribution of glia[35]. Among different types of glia, microglia have been a focal point of interest[35], because they are the major CNS resident immune cells that regulate neuroinflammation during pain pathogenesis[9]. Microglia are activated in the spinal cords of animals administered with opioids[17; 23], and implicated in morphine-induced prolonging of neuropathic pain[18]. Selective inhibition by minocycline or by anti-MAC1 (CD11B)-saporin impaired the expression of thermal hyperalgesia induced by morphine[17; 20]. Conditional knockout (CKO) of BDNF induced by CD11b-Cre and CKO of Mu opioid receptor (MOR) driven by Cx3cr1-Cre were reported to impair thermal OIH[17; 33]. It was also reported that opioids simulated microglia via Toll-like receptor 4 (TLR4) [21], and activated microglia may promote OIH expression by releasing signaling molecules such as cytokines, BDNF and ATP[35]. These lines of evidence appear consistent with the popular conception that microglia support the development of OIH and are an attractive therapeutic target for the paradoxical pain. However, glial expression of MOR remains controversial[12; 26; 29] (and refs cited therein). Studies from Corder et al. suggest that morphine acts through peripheral nociceptor but not microglia to induce OIH[12], which evoked considerable discussion in the field about the contribution of microglia to OIH[36].

In this study, we used two specific and efficient approaches to specifically ablate microglia in mice or rats. Strikingly, these approaches did not cause significant impairment of mechanical and thermal hyperalgesia induced by repeated administration of morphine. Our data provide direct evidence to show that microglia are dispensable for OIH development.

## Results

As microglia probably modulate pain pathogenesis specifically in males but not in females[30; 37], we first performed experiments with C57BL/6 male mice. We repeatedly administered the animals with morphine (20 mg/kg/day for 4 days; i.p.), and monitored the temporal profile of the expression of mechanical hyperalgesia by von Frey tests. We observed progressive expression of mechanical hyperalgesia after the first day of morphine administration, which peaked at day 3 or 4 (Fig. 1A). The hyperalgesia remained even after morphine administration stopped (Fig. 1A).

**FIGURE 1.**
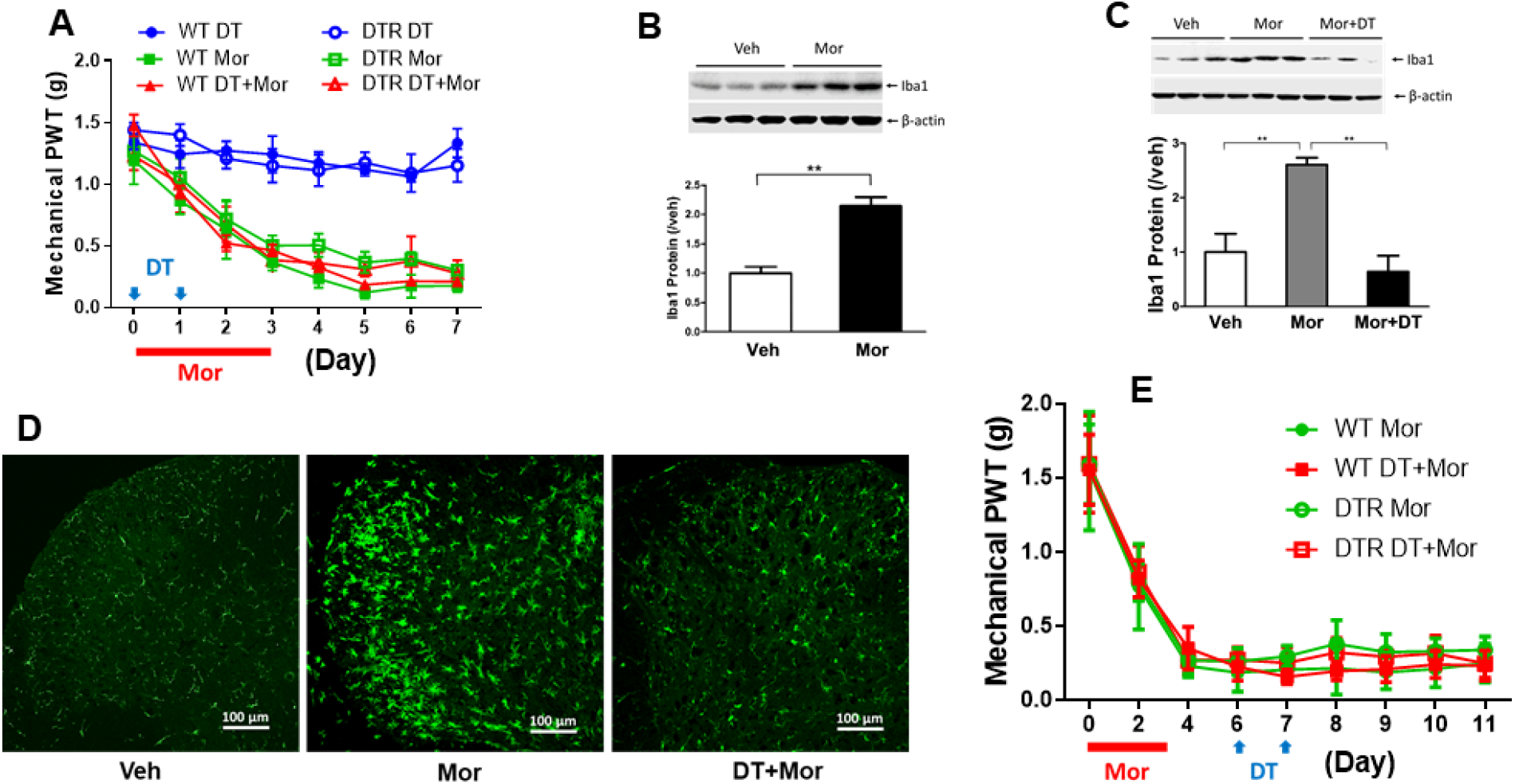
Microglial ablation in male mice by a genetic approach has no effect on development and maintenance of mechanical hyperalgesia induced by morphine. A. Effect of early DT administration on mechanical sensitivity of WT and CD11b-DTR transgenic mice with or without morphine (Mor) treatment. DT was injected (i.t.; indicated by blue arrows) at the beginning of morphine administration (i.p. daily for 4 days, indicated by the red bar) to evaluate the role of microglia in OIH expression. Von Frey tests were performed to measure the hind paw withdrawal threshold (PWT). Von Frey testing was performed 2 hours prior to morphine (i.p.) and DT (i.t.) treatment, to avoid the interference of potential acute analgesic effects of morphine administration. DT treatment of the DTR transgenic mice (empty red triangle) did not cause detectable effect on the expression of morphine-induced hypersensitivity compared with the OIH profiles of various control groups, including morphine-treated WT (solid green square) and DTR transgenic mice without DT (empty green square). N=6 mice/group; p>0.05 (DTR+DT+Mor vs. DTR+Mor, DTR+DT+Mor vs. WT+DT+Mor, or DTR+DT+Mor vs. WT+Mor); B. Immunoblotting analysis of Iba1 protein in the spinal cord of morphine-treated WT mice at day 7 indicated in paradigm in Fig.1A (left) and quantitative summary (right) (**, p<0.01). C. Immunoblotting analysis of Iba1 protein in the spinal cord of CD11b-DTR transgenic mice at day 7, treated by vehicle (Veh), morphine (Mor) or Mor+DT (left) and quantitative summary (right; **, p <0.01). D. Immunofluorescent images of microglia in the spinal cord dorsal horn of the CD11b-DTR transgenic mice at day 7, treated by vehicle (Veh), morphine (Mor) or Mor+DT. E. Effect of late DT administration on mechanical sensitivity of WT and CD11b-DTR transgenic mice with morphine (Mor) treatment. DT was injected (i.t.; indicated by blue arrows) after the completion of morphine administration (i.p. daily for 4 days, indicated by the red bar) to evaluate the role of microglia in OIH maintenance. No detectable effects on mechanical sensitivity were observed for DT-treated DTR transgenic mice (empty red square), compared with various control groups. N=6 animals/group; p>0.05 (DTR+DT+Mor vs. DTR+Mor, DTR+DT+Mor vs. WT+DT+Mor, or DTR+DT+Mor vs. WT+Mor).

Microglial activation in response to morphine treatment was evaluated by measuring the protein level of microglial marker Iba1. Similar to previous studies[13; 17; 32; 38], we observed Iba1 increase in the spinal cord collected at day 7 (Fig. 1B). Fluorescent immunostaining of Iba1 revealed drastic increase of reactive microglia in the spinal cord dorsal horn (SDH) (Fig.1D).

To determine the role of spinal microglia in the expression of OIH induced by morphine, we used the CD11b-diphtheria toxin receptor (DTR) transgenic mouse to ablate microglia[15]. Administration of diphtheria toxin (DT) to the transgenic mice (0.8 μg/kg/day for 2 days, i.t.) at the beginning of morphine treatment resulted in complete blockage of morphine-induced increase of Iba1 protein (Fig. 1C) and microglial cells in the SDH (Fig. 1D), demonstrating that the DT administration efficiently depleted microglia.

We determined the effect of microglial ablation on the expression of mechanical hyperalgesia induced by morphine. The results showed that the transgenic mice without DT treatment developed OIH with a temporal profile similar to that of WT mice (Fig. 1A), indicating the transgene did not alter OIH expression. Interestingly, DT treatment did not cause a detectable change of OIH of the transgenic mice (Fig. 1A). Specifically, the OIH expression profile of the transgenic mice treated with DT was not significantly different from that of the transgenic mice without DT treatment, WT mice with DT treatment, or WT mice without DT treatment (Fig. 1A). These results suggest that microglial ablation in the transgenic mice does not affect the expression of mechanical hyperalgesia induced by morphine.

To assess if microglia were involved in OIH maintenance, we performed additional experiments to determine the effect of DT administered after OIH was established. We observed that injection of DT at days 6 and 7 when OIH was established, had no detectable effect on the established OIH, compared with the control groups (Fig. 1E). These data indicate that ablation of microglia in the transgenic mice does not affect the maintenance of OIH.

To validate the above observations, we determined the effect of microglial ablation using a recently developed colony-stimulating factor 1 receptor (CSF1R) inhibitor PLX5622, which specifically and efficiently depletes[16; 19; 22; 34]. Microglial depletion was reported after three days of PLX5622 administration[1], and we confirmed that almost complete ablation of mouse spinal microglia was achieved within 5 days on PLX5622 diets (data not shown). To evaluate microglial contribution to OIH expression, we used a feeding paradigm with PLX5622-containing diets that started 5 days prior to morphine treatment and continued afterward. To induced OIH, mice were repeatedly administered with morphine (20 mg/kg/day for 4 days; i.p.). Spinal cords were collected after last morphine injection to measure microglial ablation. We observed nearly complete depletion of microglia, as shown by Western blotting (Fig. 2A), fluorescent immunostaining (Fig. 2B), or flow cytometry (Fig.2C). We performed von Frey tests, and observed no evident differences between the temporal profiles of mechanical OIH expressed by the WT mice (C57Bl/6) with or without PLX5622 treatment (Fig. 2D). To assess the potential role of microglia in OIH maintenance, we administered PLX5622 after mechanical OIH was established. Von Frey tests showed that PLX5622 had no detectable effect on mechanical OIH (Fig. 2E). Thus, the data suggest that PLX5622, although it almost completely depleted microglia, did not affect both expression and maintenance of mechanical OIH.

**FIGURE 2.**
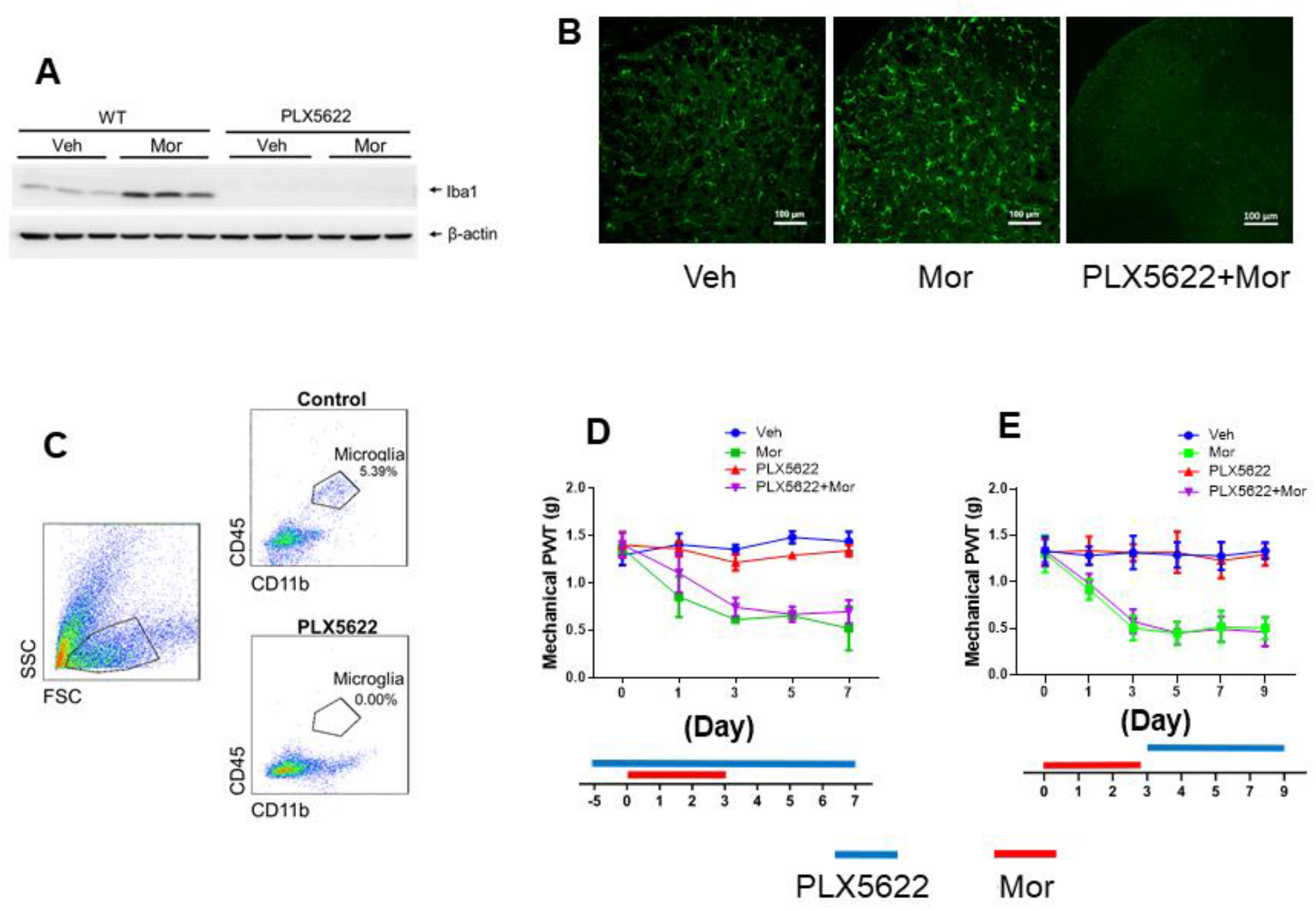
Microglial depletion in male mice by a pharmacological approach did not affect mechanical hyperalgesia induced by morphine. A-C. Microglial depletion in the spinal cord verified by immunoblotting (A), immunostaining (B) and flow cytometry analysis (C). A feeding paradigm with PLX5622-containing diets for mice (C57BL/6 males) was started 5 days prior to morphine treatment (one i.p./day for 4 days), and continued afterward. Spinal cords were collected from mice that were sacrificed at day 3 after last morphine injection, for immunoblotting (A, Iba1), immunostaining (B, Iba1) and flow cytometry analysis (C, microglia are identified as CD45-high and CD11b-high cells). All three approaches confirm the almost complete depletion of microglia in the spinal cord and the cortex (not shown), as indicated by the barely detectable Iba1 on Western blots (A), few of Iba1 positive cells on fluorescent images (B) and lack of microglia revealed by flow cytometry (C). D. Effect of microglial depletion on the expression of mechanical hyperalgesia induced by morphine in C57BL/6 males. The periods of drug administration are shown beneath the graph, with morphine administration (one i.p./day for 4 days) indicated by a red bar and PLX5622 by a blue bar (The same illustrations are used in D-E). Von Frey testing was performed to measure mechanical hyperalgesia induced by morphine treatment. PLX5622-mediated microglial depletion did not affect the mechanical OIH development. N=6 male mice/group. E. Effect of microglial depletion on the maintenance of mechanical hyperalgesia induced by morphine in C57BL/6 males. Mice were administered with PLX5622-containing diets after the completion of morphine administration period and the establishment of mechanical OIH. Von Frey tests revealed that the PLX5622 administration did not affect maintenance of morphine-induced mechanical hypersensitivity. N=6 mice/group; p>0.05 for PLX6522+Mor vs. Mor and Veh vs. PLX6522 in Fig. 2D-2E.

We also measured the potential effect of microglial ablation on the expression of thermal OIH, and found no detectable differences between the groups with or without PLX6522 administration (Fig. 3). The data suggest collectively that microglia are dispensable for the expression of both mechanical and thermal hyperalgesia induced by repetitive morphine treatment (Fig. 2; Fig. 3).

**FIGURE 3.**
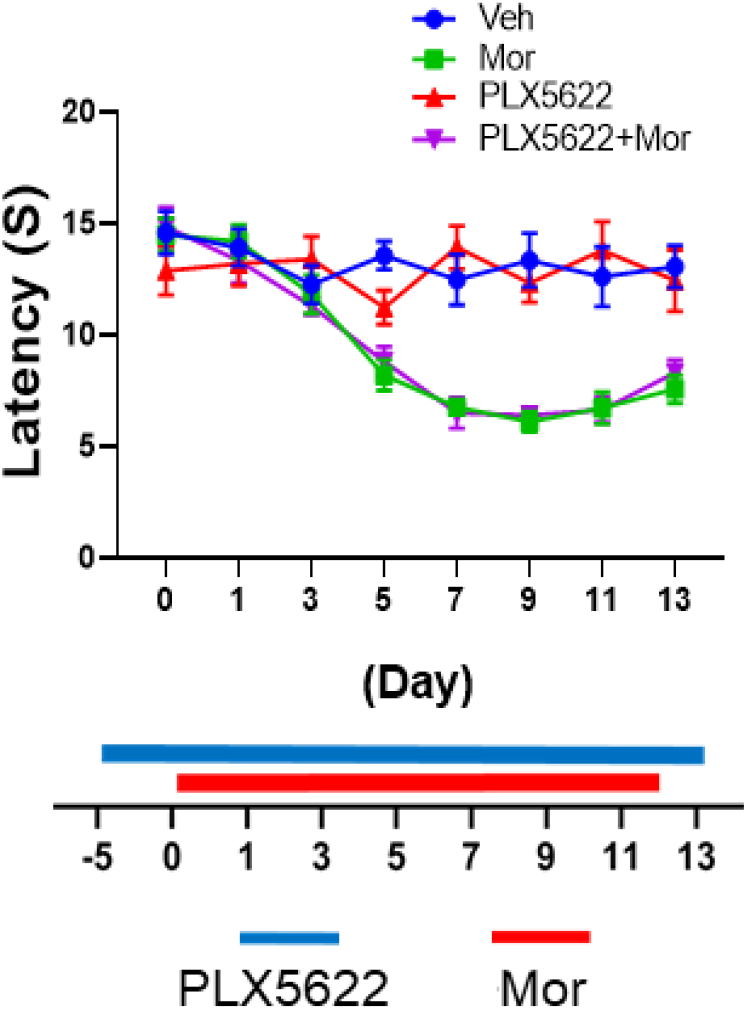
Microglial depletion in C57BL/6 maleS by a pharmacological approach did not affect thermal hyperalgesia induced by morphine. Hot plate tests were performed to measure the latency of hindpaw licking and/or shaking, or jumping. PLX5622-mediated microglial depletion did not affect the thermal OIH development. N=6 mice/group; p>0.05 for PLX6522+Mor vs. Mor and Veh vs. PLX6522.

Recent evidence suggests a sex dimorphism of microglial contribution to the pathogenesis of neuropathic pain, with a key role of microglia in male but not female rodents [30; 37]. However, the above results indicate that microglia in male mice are dispensable for the development of hyperalgesia induced by morphine. This surprising finding prompted us to test the effect of microglial ablation in females. We found that PLX5622 treatment did not cause detectable impairment of the mechanical OIH in females either (Fig. 4), indicating microglia are dispensable for mechanical OIH in both males and females.

**FIGURE 4.**
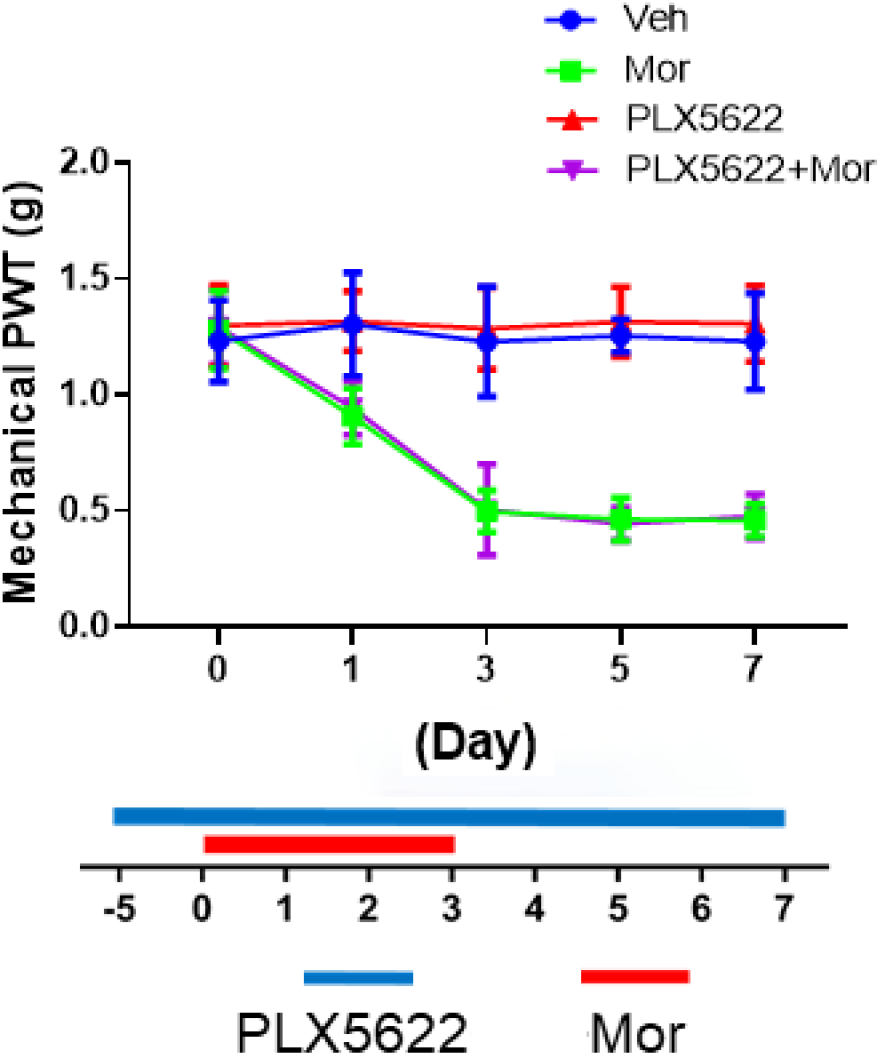
Microglial depletion in C57BL/6 females did not affect on the expression of mechanical hyperalgesia induced by morphine. Von Frey tests showed that the PLX5622 administration did not affect the expression of morphine-induced mechanical hypersensitivity. N=6 mice/group; p>0.05 for PLX6522+Mor vs. Mor and Veh vs. PLX6522.

We also tested the effect of PLX6522 on rats (male, Sprague-Dawley), and observed that PLX5622 administration completely reversed microglial increase induced by morphine although the ablation was not as complete as in mice (Fig.5A). Von Frey tests showed no detectable effect of PLX6522 on mechanical OIH induced in rats (Fig.5B).

**FIGURE 5.**
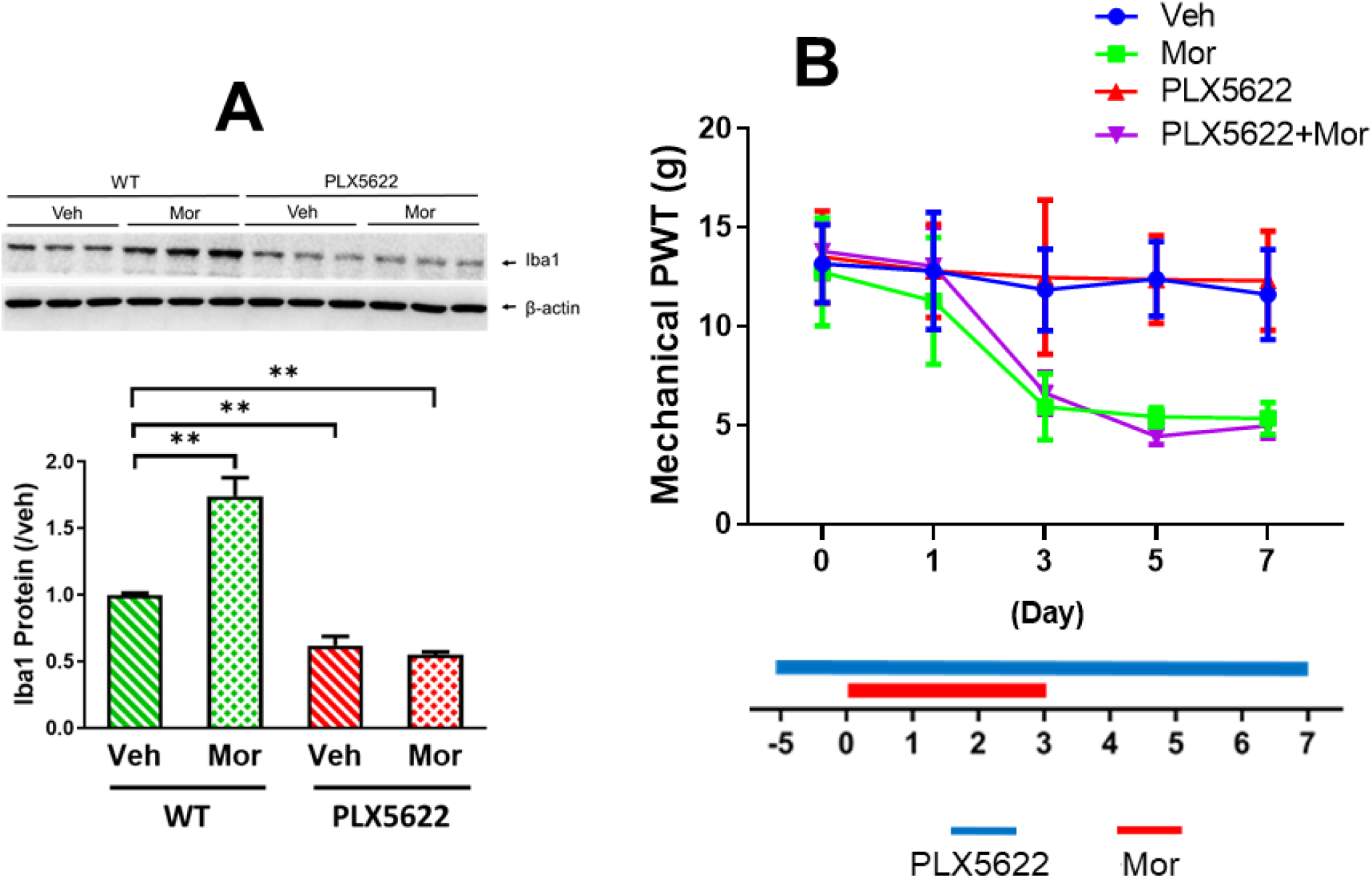
Microglial ablation by PLX6522 did not affect mechanical hyperalgesia in rats. A. PLX6522-mediated microglial ablatin in rat spinal cords. Male rats (Sprague-Dawley, 180-200g) were fed with PLX6522-containing diets for 5 days before the start (day 0) of morphine administration (20mg/kg; i.p./daily for 4 days). Spinal cords were collected at day 7. Marked decrease of spinal microglia (as indicated by Iba1 levels) was induced by PLX6522. B. Effect of PLX6522-mediated microglial ablation on OIH expression in male rats. PLX5622 administration did not affect the development of morphine-induced mechnical hypersensitivity in rats, measured by Von Frey tests. N=6 male rats/group; p>0.05 for PLX6522+Mor vs. Mor and Veh vs. PLX6522.

When we compared the results of tests on mechanical nociceptive sensitivity between control male mice received sham treatment with vehicles and the corresponding groups treated only for ablating microglia, by either genetic (Fig. 1A) or pharmacological approaches (Fig. 2D; Fig. 2E), we observed no detectable differences in nociceptive responses between the two groups (Fig. 1A; Fig. 2D; Fig. 2E). Similar results were observed for female mice (Fig. 4) and rats (Fig. 5B). We did not observe difference of thermal nociceptive sensitivity between the control and PLX6522-treated mice neither (Fig. 3). Collectively, these data suggest microglial ablation did not affect the basal nociceptive responses.

## Discussion

We are interested in interrogating the potential role of microglia in OIH, by determine the effect of microglial ablation by different approaches, in different genders, and different rodent species (mouse and rat). We observed that microglial ablation introduced during the induction of mechanical OIH did not affect OIH expression in male mice (Fig. 1A; Fig. 2D). Similarly, we observe no detectable changes of mechanical OIH expression in female mice with microglial ablation (Fig. 4). When ablating microglia was performed after mechanical OIH had been established, we did not observe changes of OIH maintenance (Fig. 1E; Fig. 2E). Furthermore, microglial ablation also did not change the expression of thermal OIH (Fig. 3). Finally, microglial ablation in rats did not change the expression of mechanical OIH (Fig. 5). These findings suggest that microglia are dispensable for the expression and maintenance of OIH. Our data are consistent with recent randomized studies with both laboratory animals and human patients on the effect of the glial inhibitor minocycline[2; 4].

Our data superficially seem to be inconsistent with some of the previous reports that show the critical roles of microglia in opioid-induced hyperalgesia[17; 20]. Because the expression of OIH is modulated by many molecular and cellular processes[35], it is well possible that studies from different laboratories may be performed under different conditions that differentially affected these processes. Different studies may differ in the animals used (ages, genders and species), the forms and doses of opioids, and/or the routes or paradigms of drug administration. These types of experimental manipulations may modify the pathogenic processes of OIH that are differentially contributed by microglia.

Microglia are only required for the development of mechanical pain hypersensitivity induced by nerve injury in male but not female mice and rats[30; 37]. This sex dimorphism of microglia raises the interesting possibility that mechanical pain hypersensitivity induced by different pathogenic causes in male mice may differentially depend on microglia. In this context, it is important to note that both males and female mice were used in this study. Our data suggest that microglia are dispensable in both males and female mice for the development of mechanical hypersensitivity induced by morphine.

Different approaches of microglial manipulation may contribute to the observed discrepancy. For instance, non-specific effects of saporin-mediated cell ablation on astrocytes were reported[28], which may complicate result interpretation when saporin-based approaches are used to ablate microglia. Hence, it is critical to use complementary strategies of microglial manipulations. CD11b-Cre-driven CKO of BDNF and Cx3cr1-Cre-driven CKO of MOR were reported to impair thermal OIH [17; 33]. However, because CD11b and Cx3cr1 are also expressed in multiple other cell types in addition to microglia [24; 27; 31], the possibility that the CKOs in non-microglial cells caused the observed impairment of thermal OIH in mutant mice has not been conclusively excluded.

In conclusion, we have used two different approaches to ablate microglia. Our results show that the development of OIH induced by morphine is not affected even after most of microglia are depleted by PLX5622 (Fig. 2A-2C). Both mechanical and thermal OIH can still be induced by morphine, with no detectable impairment, when microglia are ablated. Both male and female mice can still develop OIH, with no detectable defect, after microglia ablation. In addition, similar to mice, rats with microglial ablation also still express OIH as controls. These findings strongly suggest that microglia are not essential for the development of OIH.

## Materials & Methods

### Animals

Adult male or female C57BL/6 mice (8–10 weeks) and CD11b-DTR (006000; B6. FVB-Tg (ITGAM-DTR/EGFP)34Lan/J) mice were purchased from The Jackson Laboratory (Bar Harbor, ME, USA). The expression of diphtheria toxin receptor in CD11b-DRT mice is controlled by the human ITGAM (integrin alpha M) promoter (CD11b), and infusion of diphtheria toxin induces the reversible depletion of microglia/macrophages in the transgenic mice[15]. Additionally, adult male, Sprague-Dawley rats (180–200 g) were also used. Animal procedures were performed following protocols approved by the Institutional Animal Care and Use Committee of the University of Texas Medical Branch.

### Materials

Morphine sulfate was purchased from West-Ward (Eatontown, NJ, USA). PLX5622-containing rodent diet (each kilogram contains 1200 mg PLX5622) was purchased from Research Diets (New Brunswick, NJ, USA). Diphtheria toxin (DT) from Sigma; HibernateA and HibernateA-Ca from BrainBits (Springfield, IL, USA); papain from Worthington Biochemical (Lakewood, NJ, USA); Optiprep Density Gradient Medium from Sigma (St. Louis, MO, USA). Antibodies used for immunoblotting were: anti-IBa1 (1:1000, Wako: 016-20001); anti-GFAP (1:1000, Millipore: MAB360) and anti-β-actin (1:1000, Santa Cruz Biotechnology: sc-1616-R). The antibody for immunohistochemistry was anti-IBa1 (1:200, Wako: 016-20001). Antibodies used for flow cytometry were: APC-anti-CD11b (Biolegend: 101211), FITC-anti-CD45 (Biolegend: 103107) and anti-CD16/32 (Biolegend: 101302).

### Drug administration

In this study, animals were intraperitoneally (i.p.) injected with morphine sulfate (20 mg/kg) daily for 4 consecutive days to lead to and maintain hyperalgesia. For investigating the effect of microglia on the development of morphine-induced hyperalgesia, DT (20 ng) was intrathecally (i.t.) injected to CD11b-DRT or wild type mice daily for 2 days. To establish the PLX5622-induced microglia ablation model, C57BL/6 mice or SD rats were fed with rodent diets containing PLX5622 for 5 days prior to administration of morphine. For identifying the influence of microglia on the maintenance of morphine-induced hyperalgesia (OIH), the DT injection schedule was delayed until the 7th day after the first administration of morphine in CD11b-DRT or wild type mice; in the PLX5622-induced model, mice were not fed with PLX5622-containing diets until after the last injection of morphine. Mice treated with vehicle were used as controls.

### Measurement of mechanical and thermal nociception

Mechanical nociceptive hypersensitivity in mice was measured as described[6]. The plantar surface of the hind paw of mice in the resting state were stimulated by the calibrated von Frey filaments (Stoelting, Wood Dale, IL, USA) and paw withdrawal thresholds (PWT) were recorded using the Dixon up and down paradigm. In rat case, PWT were assessed on the ventral surface of the hind paw using von Frey filaments (Stoelting, Wood Dale, IL, USA) and the up-down method[8; 14; 40]. For thermal sensitivity test, paw withdrawal latency to a thermal stimulus was measured using a solid state laser method as described[7]. The latency to hindpaw licking and/or shaking, or jumping were recorded. All tests were conducted 2 hours before treatment by an experimenter who did not know the details on the treatments.

### Spinal cell dissociation

HibernateA (HABG) media containing 30mL HibernateA, 600μL, 88μL 0.5mM Glutamax and 2μL of 1% penicillin and streptomycin was prepared prior to dissection. Papain mixture containing 12mg papain, 6 ml HibernateA-Ca media and 15μL 0.5mM Glutamax was placed in a 37°C water bath for 20 minutes and then placed on ice until use. Animals were anesthetized and cardiac perfusion was carried out followed by rapidly dissecting lumber segment of spinal cord. Tissues were placed in 6ml HABG on ice and cut into 0.5mm pieces before transferred into 6ml papain mixture. The mixture with tissues was incubated in 37°C for 30 minutes with occasional shaking. After incubation, tissues were transferred to 2ml HABG as little papain as possible and triturated 15-20 times using polished glass Pasteur pipettes avoiding bubbles followed by settling in room temperature for 1 minute. The supernatant was transferred to an empty 15ml tube. 2ml HABG was added to the sediment and two times of repeats for triturating, settling and transferring supernatant were carried out and finally got 6ml cell suspension. To separate the debris from the cell suspension, a four layers of density gradient media was created as described[5]. Cell suspension was carefully pipetted on top of the density gradient with little mixing. The density gradient was centrifuged at 800g for 15 minutes at 22°C and then the top layer was aspirated and discarded. Another 5ml HABG was added to dilute the density gradient and centrifuged for 10 minutes at 1000rmp at 22°C. After centrifuging, supernatant was discarded and 2ml HABG was added to resuspend the cell pellet.

### Flow cytometry staining

Dissociated spinal cells were mixed with FACS buffer containing 1xPBS, 5%FBS and 2mM EDTA and centrifuged at 800g for 5 minutes at 4°C. Cell pellet was resuspended by FACS buffer and then counted to adjust cell number to a concentration of 2×10^6^ cell/mL with FACS buffer. Cell suspension was centrifuged at 400g for 8 minutes in 4°C and then resuspended by 100μL FACS buffer containing 1μL Fc blocker and incubated for 5 minutes avoiding light. Antibodies were added to cell suspension and mixed well followed by incubating at 4°C for 30 minutes. After incubation, samples were washed using FACS buffer twice and centrifuged at 400g for 8 minutes in 4°C and resuspended in 300μL FACS buffer for flow test.

### Western blotting analysis

Mice were anesthetized with 3% isoflurane and sacrificed for collecting the L4-L6 lumbar spinal cord segments. Tissues were homogenized in RIPA lysis buffer (1% Nonidet P-40, 50 mM Tris–HCl pH 7.4, 150 mM NaCl, 0.5% sodium deoxycholate,, and 1 mM EDTA pH 8.0) with a protease inhibitor cocktail (Sigma). BCA Protein Assay kits (Thermo) were used to determine protein concentrations. Equal amounts of protein were loaded and separated by 12% SDS-PAGE and transferred to polyvinylidene fluoride membranes. The membranes were blocked in 5% nonfat milk in TBST buffer for 1 hour at room temperature and then incubated with anti-IBa1 (1:1000, Wako: 016-20001); anti-GFAP (1:1000, Millipore: MAB360) and anti-β-actin (1:1000, Santa Cruz Biotechnology: sc-1616-R) in TBST buffer overnight at 4°C. After washing with TBST buffer, membranes were incubated with HRP-conjugated secondary antibody. Enhanced Chemiluminescence kits (Pierce) were used to visualize protein bands, and NIH ImageJ software was used for quantification. β-actin was used here as a reference control.

### Immunohistochemistry

Mice were anesthetized with 3% isoflurane and transcardially perfused first with 20 ml of 0.01 M PBS and then with 30 ml of 4% paraformaldehyde in 0.01 M phosphate buffer. The L4 and L5 lumbar spinal cord sections were dissected out and fixed in 4% PFA solution for 12 hours at 4°C, dehydrated with 30% sucrose solution in PBS for 24 hours at 4°C, and frozen in optimal cutting temperature medium (Tissue-Tek). Tissues were cut into 15 μm sections on a cryostat (Leica CM 1900) and mounted onto Superfrost Plus microscope slides. The sections were next incubated in blocking solution containing 5% BSA and 0.3% Triton X-100 in 0.01 M PBS for 1 h at room temperature and then in rabbit anti-IBa1 (1:200, Wako: 016-20001) overnight. Sections were washed with 0.01 M PBS and incubated in FITC-conjugated secondary antibody (1:200, Jackson ImmunoResearch Laboratories). They were then stained with DAPI (Sigma). IgG from the same animal source was used as a negative control for immunostaining. Images were collected with a laser confocal microscope (Zeiss).

### Statistical analysis

Statistical analysis was conducted with Prism 5 (GraphPad) software. Data were represented as mean ± SEM from three independent experiments. One-way ANOVA was used for immunoblotting data and two-way ANOVA with a Bonferroni post hoc test was performed to analyze animal pain behavior.

## Acknowledgement

We thank Plexxikon Inc. for the use of PLX5622. SJT was supported by NIH grants R01NS079166, R01DA036165, R01NS095747 and R01DA050530, and by the Cecil H. and Ida M. Green Distinguished Chair in Neuroscience and Cell Biology. QY was supported by NIH grant R01CA208765.

## Author Contribution

XL, BLL, QY and XZ performed experiments and data analysis. SJT designed the study. SJT and XL wrote the paper.

